# Automatic identification of relevant genes from low-dimensional embeddings of single cell RNAseq data

**DOI:** 10.1101/2020.03.21.000398

**Authors:** Philipp Angerer, David S. Fischer, Fabian J. Theis, Antonio Scialdone, Carsten Marr

## Abstract

Dimensionality reduction is a key step in the analysis of single-cell RNA sequencing data and produces a low-dimensional embedding for visualization and as a calculation base for downstream analysis. Nonlinear techniques are most suitable to handle the intrinsic complexity of large, heterogeneous single cell data. With no linear relation between genes and embedding however, there is no way to extract the identity of genes most relevant for any cell’s position in the low-dimensional embedding, and thus the underlying process.

In this paper, we introduce the concepts of global and local gene relevance to compute an equivalent of principal component analysis loadings for non-linear low-dimensional embeddings. While *global gene relevance* identifies drivers of the overall embedding, *local gene relevance* singles out genes that change in small, possibly rare subsets of cells. We apply our method to single-cell RNAseq datasets from different experimental protocols and to different low dimensional embedding techniques, shows our method’s versatility to identify key genes for a variety of biological processes.

To ensure reproducibility and ease of use, our method is released as part of destiny 3.0, a popular R package for building diffusion maps from single-cell transcriptomic data. It is readily available through Bioconductor.

## 1 Introduction

Single cell RNA sequencing (scRNAseq) has massively improved the resolution developmental trajectories *Baron et al.* (2016) and allowed unprecedented insights into the heterogeneity of complex tissues Vento-Tormo *et al.* (2018); Tritschler *et al.* (2017). On the flip side, new challenges have arisen due to the amount of data that needs to be processed Angerer *et al.* (2017), higher levels of technical and biological noise *Yuan et al.* (2017), and identification and interpretation of known and novel cell types Pliner *et al.* (2019). To exploit the new opportunities and deal with the new challenges, a large number of algorithms and tools have been developed Zappia *et al.* (2018).

Dimension reduction methods create a low dimensional embedding of the high dimensional gene expression space and are widely used. Such embeddings serve as a visual overview of the data on which gene expression profiles and per-cell or per-cluster statistics can be compared. Embeddings can also serve as inputs for further downstream computational analysis. E.g., principal component analysis (PCA) is a popular technique to identify orthogonal linear combinations of genes that explain variance in the data. PCA loadings quantify the contribution of genes to each principal component and help to understand the genetic drivers of the underlying molecular processes. However, linear methods are often not able to capture the complexity of high-dimensional datasets *Haghverdi et al.* (2015), which is why nonlinear dimension reduction methods (see e.g. t-SNE Husnain *et al.* (2019), diffusion maps Husnain *et al.* (2019); *Coifman et al.* (2005); Haghverdi *et al.* (2015), UMAP McInnes *et al.* (2018); *Becht et al.* (2018), and graph-based methods Islam *et al.* (2011)) have become the standard for scRNAseq data analysis. For non-linear embeddings however, no intrinsic measure of individual genes’ contribution to each embedding dimension exists. Without such a measure, the identification of genes that drive the variability in the data requires tedious manual inspection and exclusive prior knowledge about possible target genes.

Here, we introduce *gene relevance*, a measure for a gene’s contribution to variance in low dimensional embeddings, and present a method to infer a local as well as a global gene relevance score from any kind of low-dimensional embedding. To demonstrate the utility of the method, we apply gene relevance to several datasets. In a blood cell dataset from mouse embryos, we are able to automatically identify genes involved in embryonic blood differentiation. Gene relevance is available as part of the R package *destiny* Angerer *et al.* (2016).

## 2 Results

We define gene relevance as a measure of how much a gene contributes to the cell-to-cell variability in a low dimensional embedding of a scRNAseq dataset as a function of this embedding (see Fig. 1a-c). It can be interpreted as a generalization of PCA loadings to non-linear dimensionality reduction techniques. Note that PCA loadings are constant with respect to the PC space while feature importance in a non-linear embedding is naturally a non-constant function of this embedding. A ranking of genes based on their relevance is built for every cell of the embedding. These rankings, then, can be combined to obtain a measurement of the “local” or “global” relevance of each gene (see Fig. 1d-e and **Methods**), which highlight genes relevant in small or large cell subpopulations, respectively. To explore and visualize the results further, the method also provides a “gene relevance map”, where the locally most relevant genes are displayed along with the corresponding region of the embedding (see Fig. 1f).

**Fig. 1.**
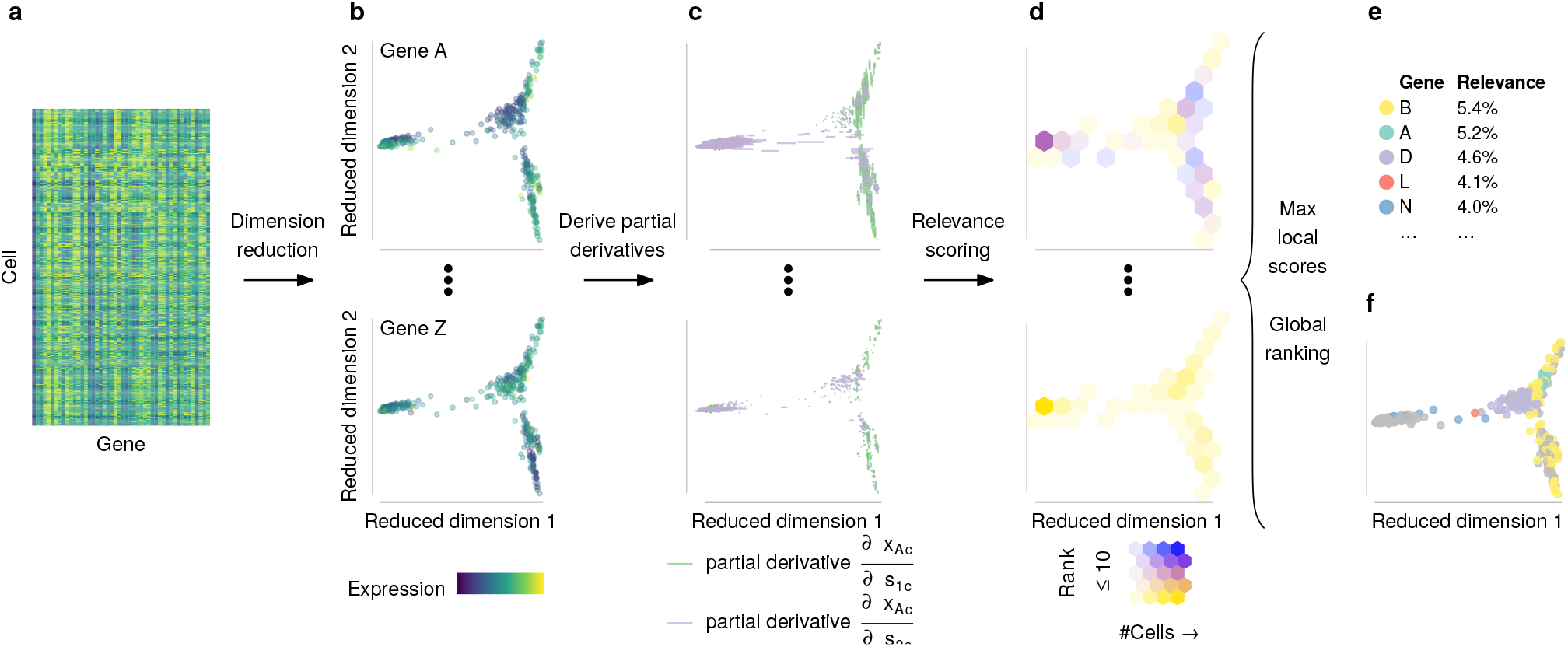
The gene relevance concept. A gene expression matrix (a) from a single cell RNA sequencing experiment is reduced to a low-dimensional embedding (b), with each dot representing a cell, and the color representing the expression of gene A, B, …, Z. Expression changes are calculated from estimates of partial derivatives with respect to the embedding (c), which results in one value per cell×gene×dimension combination. We score the relevance of each gene in each cell according to the partial derivatives’ F1 norm. This score indicates how locally relevant each gene is (d). The fraction of cells ranking a given gene above a threshold defines a global gene relevance score. In our illustrative example, gene B has been ranked among the top 10 genes in 5.4% of all cells (e). To identify the relevant genes for a particular local process in the embedding, the local scores are smoothed before the gene with the highest local score is selected (f).

We demonstrate our method on a scRNAseq dataset of blood progenitors and blood cells from mouse embryos Scialdone *et al.* (2016) (see Fig. 2a and **Methods** for more details). In the original publication, this data was used to reconstruct a trajectory representing primitive erythropoiesis, along which blood marker expression increases and other markers (such as endothelial cells) decrease. There, an ad-hoc method was devised to find important genes in the 2D diffusion map embedding of the data. Here we show how our method can be used “out of the box” to rank genes based on their local and global relevance.

**Fig. 2.**
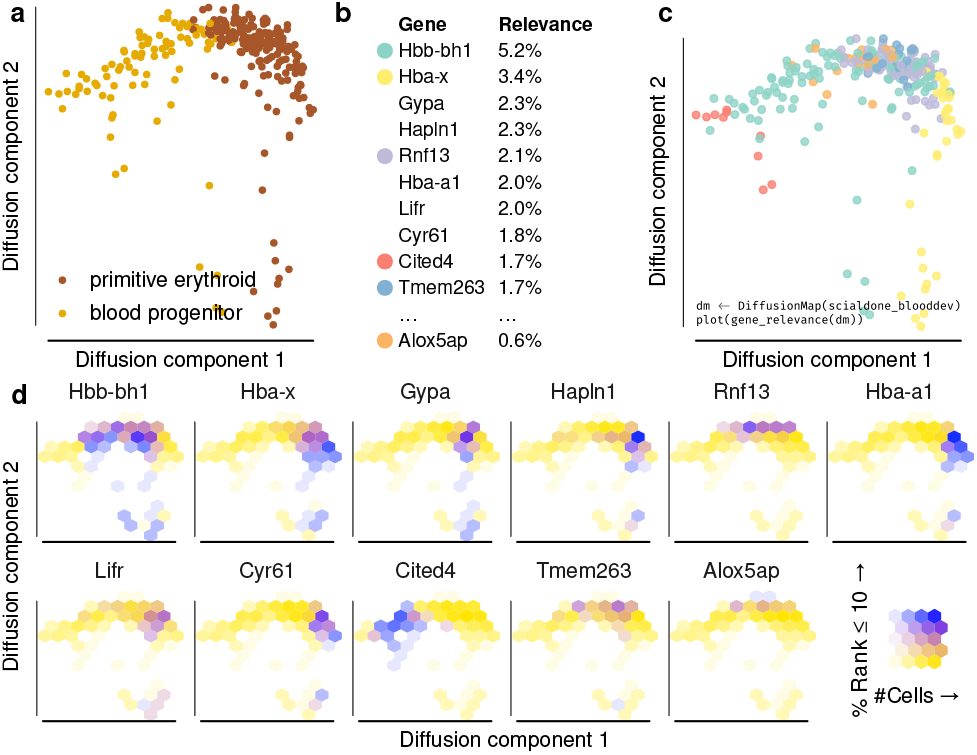
Gene relevance automatically detects drivers of embryonic blood development. (a) Diffusion map of 271 single hematopoietic progenitor cells from mostly day 7.5 and 7.75 mouse embryos, profiled in Scialdone et al. (2016) (b) Global gene relevance identifies Hbb-bh1 and Hba-x as genes that change most dramatically during hematopoietic development. (c) Local gene relevance in the diffusion space reveals the contribution of relevant genes in specific regions of the process. The genes corresponding to each color are shown in panel b. (d) Gene relevance maps detail the areas where the contribution of genes is highest. Alox5ap shows a high local relevance in the top region of the diffusion map and has been implicated with early blood development Ibarra-Soria et al. (2018).

First, we ranked all highly variable genes according to their global gene relevance (see Fig. 2b). As expected, the high-ranking genes are mostly associated with blood development, including the hemoglobin genes Hba-a1, Hba-x, Hbb-bh1, and the erythrocyte membrane genes Gypa and Cited4 Yahata *et al.* (2002). The genes Cyr61 and Hapln1 are involved in extracellular matrix and important for development of the cardiovascular system Latinkić *et al.* (2001). The top of the list has a good overlap with the ad-hoc method in Scialdone *et al.* (2016): 4 genes are shared between the top ten of both lists, and we find a Rank-Biased Overlap of *RBOp* = 0.48, where we used *p* = 0.9, which assigns ~86% of the weight to the first 10 genes Webber *et al.* (2010).

Second, we created a local gene relevance map (Fig. 2c). Five out of the six locally most relevant genes (see Fig. 2c) are among the ten most globally relevant ones. Interestingly, Alox5ap is included only in the local gene relevance map, because its contribution is confined to a small region of the diffusion space (bottom right panel in Fig. 2d) and hard to detect at the level of gene expression (see Suppl. Fig. 1). This gene was not discovered by the ad-hoc method of Scialdone *et al.* (2016), but it has been recently found to be important in early blood development Ibarra-Soria *et al.* (2018). Locally and globally relevant genes can also be inferred in other embeddings such as t-SNE Maaten and Hinton (2008) and UMAP *Becht et al.* (2018), with a high overlap of relevant genes (*RBO*(*p* = 0.9) = 0.53, 4 of the top ten relevant genes are identical. See Suppl. Fig. 2).

Applied to other scRNAseq data sets, we showcase versatility and ease of application of our method. In a data set of human endocrine cells Veres *et al.* (2019), gene relevance maps detect genes driving the separation of subpopulations in the embedding (see Suppl. Fig. 3a), in accordance to the markers identified in the original paper. In human brain organoid cells Gray Camp *et al.* (2015), we detect relevant genes different from the markers specified in the paper. The reason seems to be a low density region between mesenchymal cells and neurons/neural progenitors (see Suppl. Fig. 3b). The found genes therefore seem to mostly drive the difference between progenitors and neurons: TXNRD1 plays a vital role for neuron progenitor cells Soerensen *et al.* (2008), the selenoprotein SELT protects neurons against oxidative stress in mouse models Boukhzar *et al.* (2016), and CRABP1 modulates the neuronal cell cycle in mice Lin *et al.* (2017). Finally, we applied gene relevance to mouse embryonic stem cells grown in different culture media Kolodziejczyk *et al.* (2015). As expected for cells in a relatively homogenous pluripotent steady state, the relevant genes were enriched for cell cycle and other housekeeping gene ontology processes (see Suppl. Fig. 5).

As a sanity check, we apply gene relevance to scRNAseq data from embryonic stem cells cultured in three different pluripotency retaining media. We expected to find a homogenous, steady state cell population. Indeed the relevant genes for diffusion map embedding of all three media turned out to be involved in housekeeping, metabolic and proliferation pathways (see Suppl. Fig. 5).

## 3 Discussion

We presented a method that is able to reliably detect relevant genes from low dimensional embeddings of scRNAseq data. More specifically, our method computes both a global and a local gene relevance score: global gene relevance identifies the main drivers of the cell-to-cell variability in the whole embedding; local gene relevance picks up genes relevant in smaller regions of the embedding, e.g. to identify important genes in rare cell sub-populations. In addition to a gene ranking based on global relevance, the method also provides graphic tools to visualize the local gene relevance (see Fig. 1e) and the changes in gene expression levels within the embedding (see Fig. 1c and Suppl. Fig. 1). It can be used for any single cell data set and any dimensionality reduction technique.

We applied our method to three datasets, including one from mouse embryonic blood progenitors, where we show that it performs comparably to a technique custom-made for the dataset. Interestingly, our method identifies Alox5ap (Fig. 2), a gene that was recently shown to be important for blood development in a later publication Ibarra-Soria *et al.* (2018). In two other examples, we used human cells, endocrine Veres *et al.* (2019) and from brain organoids Gray Camp *et al.* (2015), showing that the method works robustly in varied conditions.

Other methods to identify important genes from scRNAseq data exist, but most of them aim to find marker genes that can best distinguish different cell types Delaney *et al.* (2019). Conversely, the method we presented is unsupervised and does not rely on cell type annotation.

Recently, two computational methods have been developed to identify variable genes in spatial RNAseq datasets, trendsceek and SpatialDE Edsgärd *et al.* (2018); Svensson *et al.* (2018). While these methods were designed to find patterns in spatial transcriptomic datasets, they can also be used to identify relevant genes in low-dimensional embeddings of scRNAseq datasets (see Suppl. Fig. 6 in *Edsgärd et al.* (2018)). We compared our approach to trendsceek and found similar genes (see Suppl. Fig. 6). Our method completed in 6.5 seconds while trendsceek needed 1080 seconds and only ran successfully on the exact data provided in its R package. SpatialDE returned a perfect score for a too large number of genes to be useful. This is probably related to both methods being optimized towards identifying spatial patterns. Moreover, neither method allows estimation of local gene relevance.

To summarize, our gene relevance method is a fast and versatile exploratory tool that can help identify the biological processes and reveal the presence of potentially rare cell sub-populations. It is available online, easily applicable, and faster than model fitting a pproaches. W hile we focussed our discussion on scRNAseq datasets, our method can be applied to virtually any kind of dataset where low-dimensional embeddings are obtained, including, for instance, single-cell epigenomic *Shema et al.* (2018) and mass cytometry data Spitzer and Nolan (2016).

## 4 Methods

### Single cell RNA sequencing data

We used count data from 271 cells mostly from the neural plate (embryonic day 7.5) and head fold (embryonic day 7.75) development stages of mouse embryos, published in Scialdone *et al.* (2016). There, the libraries were constructed using the Smart-seq2 protocol, read counts were obtained via HTseq-count *Scialdone et al.* (2016). The 271 cells we used correspond to the clusters annotated as “blood progenitor” and “primitive erythroid” in the original publication. We selected highly variable genes using the method of *Brennecke et al.* (2013) because of its stable performance Yip *et al.* (2018), and embedded the log-transformed data using the diffusion map implementation destiny Angerer *et al.* (2016).

### Neighborhoods

If a k nearest neighbor (kNN) search has been performed as part of the embedding, it can be efficiently u sed f or estimating the gene relevance. To perform the kNN search, destiny offers the choice between euclidean distance, cosine distance, and spearman rank correlation distance. The latter was used in all analyses performed for this paper.

### Local gene relevance

We define local gene relevance of gene *g* ∈ {1, …, *G*} in cell *c* ∈ {1, …, *C*}, *LR*(*gc*), as the Frobenius norm *F*(*d*_*gc*_) of the differential *d*_*gc*_:

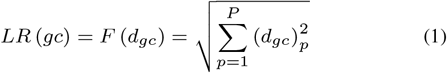

The differential *d*_*gc*_ of gene *g* in cell *c* describes the change in gene expression *x*_*gc*_ along a change in embedding coordinates *s*_*pc*_, where *p* ∈ {1, …, *P*} is the embedding dimension and *d*_*gc*_ corresponds to the partial derivatives of the gene expression with respect to each embedding coordinate:

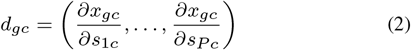

We estimated *d*_*gc*_ from the cells’ neighborhood *NN*_*k*_(*c*) in gene expression space, approximating using finite differences:

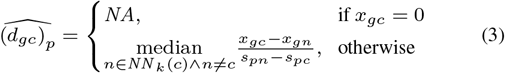

### Global gene relevance

In each cell *c*, genes can be ranked according to their local relevances LRM_*gc*_, from most to least relevant. Given the ranks 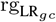 of gene *g* and a rank cutoff rg_max_, we define global gene relevance 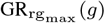 of gene *g* as:

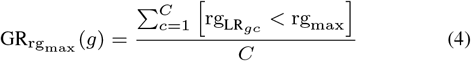

with the iverson bracket notation

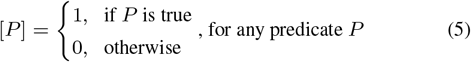

### Gene relevance maps

For a set of genes of interest Ω ∈ {1, …, *G*} (which can be chosen, e.g., among those with highest global relevance) and each cell *c*, we define the locally most relevant gene 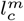 after a number of smoothing steps *m*:

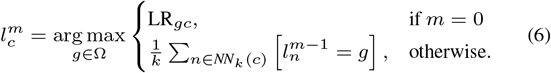

During a smoothing iteration, we replace the local gene relevance score of cell *c* and gene *g* with the fraction of neighbours that have *g* as the most relevant gene.

## 5 Availability of data and materials

The datasets analysed within this publication are available from their original publications as follows: The differentiating mouse embryonic stem cell data *Scialdone et al.* (2016) is available at http://gastrulation.stemcells.cam.ac.uk/scialdone2016, the pluripotent mouse embryonic stem cell data Kolodziejczyk *et al.* (2015) at https://www.ebi.ac.uk/teichmann-srv/espresso/, the human brain organoid data Gray Camp *et al.* (2015) at GSE75140, and the human endocrine cell data Veres *et al.* (2019) at GSE114412.

## 7 Authors’ contributions

PA designed the analysis, implemented the method. AS interpreted the results and wrote the paper with AS and CM. DF contributed to the mathematical description of the gene relevance concept. FT contributed the initial idea of gene relevance. CM supervised the study.

## Acknowledgements and Funding

D.S.F. acknowledges financial support by a German research foundation (DFG) fellowship through the Graduate School of Quantitative Biosciences Munich (QBM) (GSC 1006) and by the Joachim Herz Stiftung. FT and CM acknowledge support from the DFG funded collaborative research center SFB 1243.

