## Supplementary Materials for "Automatic identification of relevant genes from low-dimensional embeddings of single cell RNAseq data"

*Bioinformatics*
doi.10.1093/bioinformatics/xxxxxx
Advance Access Publication Date: 4th of September 2019
Original Paper

Gene Expression

### Supplementary Figures

**Angerer *et al.***

#### Abstract

Supplementary Figures for the paper *Automatic identification of relevant genes from low-dimensional embeddings of single cell RNAseq data*

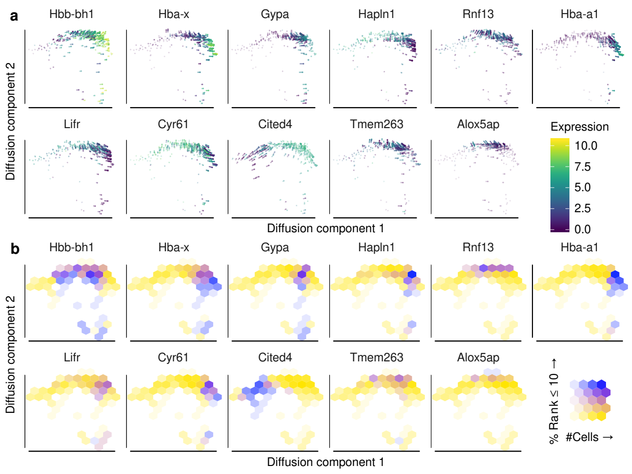


**Suppl. Fig. 1.** Gene relevance automatically identifies drivers of a developmental process. Expression levels and corresponding differentials of selected genes (a) and local gene relevance maps for the same genes (b) in a diffusion map of the embryonic blood dataset. Obvious gene expression changes can be identified from mapping the expression in single cells in a low dimensional embedding (e.g. for Cited4, Cyr61, Hba-a1), while the direction and strength (line length) of the differential simultaneously visualizes gene expression changes (a). Subtle, but also important gene expression changes are more robustly identified with our concept of gene relevance (e.g. Alox5ap, Tmem263, Rnf13) as explained in Fig. 1a (b).

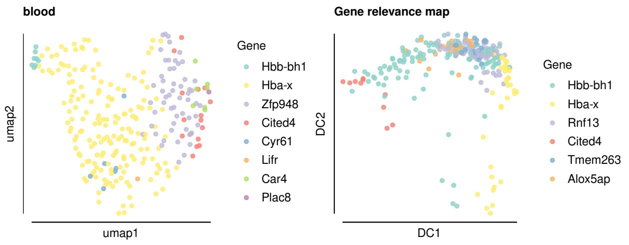


**Suppl. Fig. 2.** t-SNE (a) and diffusion map (b) embeddings of the embryonic blood dataset, with the top locally relevant genes indicated. While the embeddings have differently sized regions driven by genes their gene relevance maps share, comparable lists of genes are identified, and some patterns are conserved (e.g., Hba-x and Cited4 marking opposite ends of both embeddings).

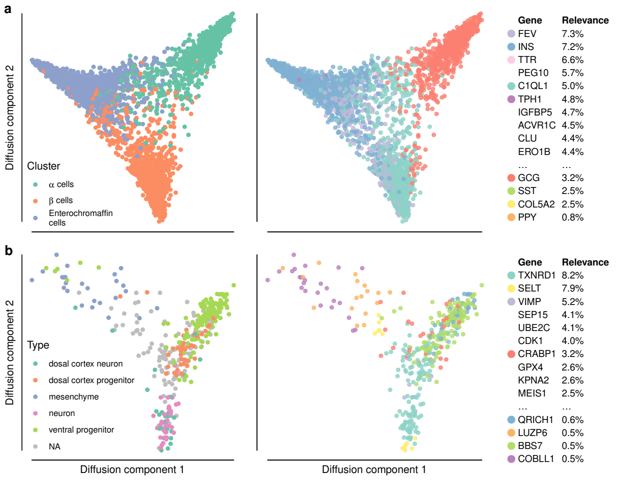


**Suppl. Fig. 3.** Gene relevance automatically identifies drivers of tissue heterogeneity and differentiation in two human scRNAseq datasets. (a) Reaggregated stem cell derived endocrine cells Veres et al. (2019). FEV, GCG, INS, and TPH1 are mentioned as markers for the respective cell types in the original paper. DLK1 was manually excluded due to its dominating of the gene relevance map. The gene relevance map reveals that FEV mostly drives the distinction between enterochromaffin (EC) and $\beta$ cells, while GCG takes a similar role for EC vs. $\alpha$ cells. (b) Embryonic stem cell derived brain organoids Gray Camp et al. (2015). The relevant genes TXNRD1, SELT, VIMP, and CRABP1 are known to play roles in neuronal proliferation and differentiation.

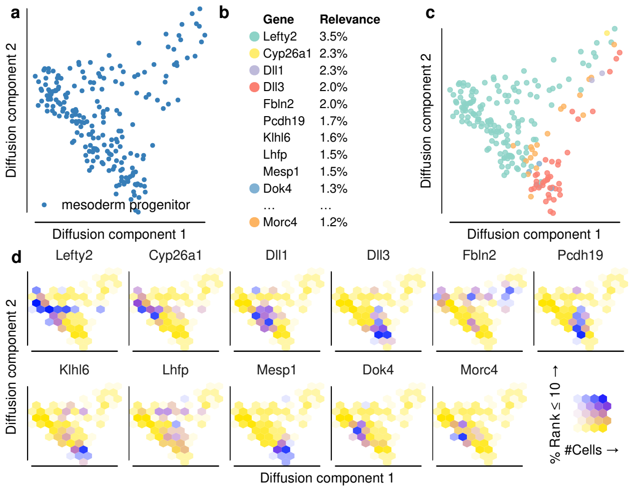


**Suppl. Fig. 4.** Gene relevance successfully identifies relevant genes in mesodermal progenitor cells. (a) Diffusion map of 216 mesodermal progenitor cells profiled in Scialdone et al. (2016). (b) List of globally most relevant genes in the diffusion map and (c) corresponding local gene relevance, with the command used to generate it. (d) Local gene relevance maps for the most locally and globally relevant genes, show relevance patterns not visible on the gene relevance map (c).

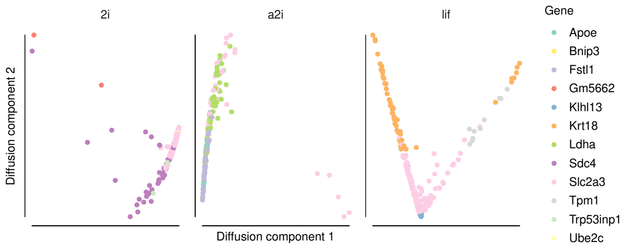


**Suppl. Fig. 5.** Diffusion maps of mouse embryonic stem cells grown in 3 different pluripotency maintaining media (2i, a2i, and Lif, indicated on top of each plot), with locally relevant genes. Data from Kolodziejczyk et al. (2015). As expected, the identified relevant genes are responsible for housekeeping and other regular cellular functions: The top 3 enriched gene ontology terms retrieved by GOrilla Eden et al. (2009) for the three media are: 2i: cell division, positive regulation of cellular protein catabolic process, negative regulation of mitotic cell cycle, a2i: response to toxic substance, lactate biosynthetic process from pyruvate, response to oxidative stress, lif: negative regulation of cellular process, negative regulation of biological process, cellular component organization.

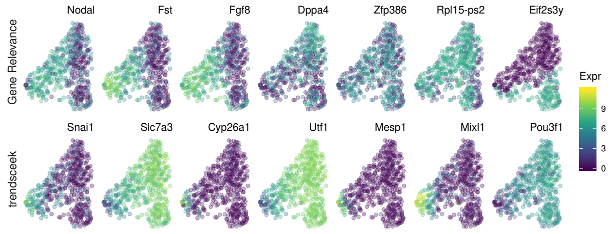


**Suppl. Fig. 6.** Trendsceek Edsgärd et al. (2018) and global gene relevance were applied to identify relevant genes from a UMAP embedding of epiblast cells from Scialdone et al. (2016). The figure shows the expression levels of the top relevant genes identified by global gene relevance (top row) and trendsceek (bottom row) overlaid to a t-SNE embedding similar to one from the trendsceek paper. Both lists include genes that play important roles in gastrulation (e.g., T, Mixl1, Fgf8, Frzb, etc), as expected for epiblast cells at this stage of development Scialdone et al. (2016).
